# Neuropeptide-dependent spike time precision and plasticity in circadian output neurons

**DOI:** 10.1101/2024.10.06.616871

**Authors:** Bryan Chong, Vipin Kumar, Dieu Linh Nguyen, Makenzie A. Hopkins, Faith S. Ferry, Lucia K. Spera, Elizabeth M. Paul, Anelise N. Hutson, Masashi Tabuchi

## Abstract

Circadian rhythms influence various physiological and behavioral processes such as sleep-wake cycles, hormone secretion, and metabolism. In *Drosophila*, an important set of circadian output neurons are called pars intercerebralis (PI) neurons, which receive input from specific clock neurons called DN1. These DN1 neurons can further be subdivided into functionally and anatomically distinctive anterior (DN1a) and posterior (DN1p) clusters. The neuropeptide diuretic hormones 31 (Dh31) and 44 (Dh44) are the insect neuropeptides known to activate PI neurons to control activity rhythms. However, the neurophysiological basis of how Dh31 and Dh44 affect circadian clock neural coding mechanisms underlying sleep in *Drosophila* is not well understood. Here, we identify Dh31/Dh44-dependent spike time precision and plasticity in PI neurons. We first find that a mixture of Dh31 and Dh44 enhanced the firing of PI neurons, compared to the application of Dh31 alone and Dh44 alone. We next find that the application of synthesized Dh31 and Dh44 affects membrane potential dynamics of PI neurons in the precise timing of the neuronal firing through their synergistic interaction, possibly mediated by calcium-activated potassium channel conductance. Further, we characterize that Dh31/Dh44 enhances postsynaptic potentials in PI neurons. Together, these results suggest multiplexed neuropeptide-dependent spike time precision and plasticity as circadian clock neural coding mechanisms underlying sleep in *Drosophila*.

## 1 INTRODUCTION

Circadian rhythms are biological processes that follow an approximately 24-hour cycle and are regulated by an internal biological clock (Brown & Schibler, 1999; Dunlap, 1999; Helfrich-Forster, 2003; Gachon *et al*., 2004; Green *et al*., 2008; Nitabach & Taghert, 2008; Buhr & Takahashi, 2013; Hardin & Panda, 2013; Dubowy & Sehgal, 2017; Cedernaes *et al*., 2019). These rhythms influence various physiological and behavioral processes such as sleep-wake cycles, hormone secretion, and metabolism (Green *et al*., 2008; Franken & Dijk, 2009; Dallmann *et al*., 2012; Cedernaes *et al*., 2019; Hill *et al*., 2020). Circadian output neurons are a group of neurons that receive input from the central circadian clock located in the suprachiasmatic nucleus of the mammalian brain and transmit timing information to different regions of the brain and body (Sun *et al*., 2001; Herzog & Schwartz, 2002; Welsh *et al*., 2010; Colwell, 2011; Mohawk & Takahashi, 2011), coordinating the circadian rhythms of various physiological processes (Hastings *et al*., 2018; Goltsev *et al*., 2022). Disruption of clock genes has been linked to sleep disorders and the development of metabolic diseases (Maury *et al*., 2014; Cedernaes *et al*., 2019; Cederroth *et al*., 2019). Conversely, aberrant nutrient signaling can affect circadian rhythms of behavior, suggesting a bidirectional interaction between circadian rhythms and metabolic processes (Huang *et al*., 2011). The circadian rhythm also interacts with metabolic processes to regulate energy balance, influencing factors such as hormone release, nutrient distribution, and basal metabolic rate (Panda, 2016). DN1 neurons are known to exhibit anatomical and functional heterogeneity, including anterior (DN1a) and posterior (DN1p) clusters. Several non-clock circuits have been identified downstream of DN1p clock neurons, including tuberculo-bulbar neurons (Guo *et al*., 2018; Lamaze *et al*., 2018) and a potentially specific class of dopaminergic neurons (Liang *et al*., 2019). Besides them, there is an important set of circadian output neurons called pars intercerebralis (PI) neurons (Cavanaugh *et al*., 2014), which receive monosynaptic inputs specifically from DN1 clock neurons (Barber *et al*., 2016), controlling the circadian regulation of sleep (Guo *et al*., 2016; Guo *et al*., 2018; Lamaze *et al*., 2018; King & Sehgal, 2020; Shafer & Keene, 2021; Tabuchi *et al*., 2021). The dynamics of the *Drosophila* circadian network involve many types of neuropeptidergic signaling (Schlichting *et al*., 2016; King *et al*., 2017; Klose *et al*., 2021), such as PDF, pigment-dispersing factor (Mertens *et al*., 2005), leucokinin (Cavey *et al*., 2016), short neuropeptide F (Shang *et al*., 2013), ion transport peptide (Hermann-Luibl *et al*., 2014), and Allatostatin A (Chen *et al*., 2016), in addition to fast amino acid neurotransmitters such as glutamate (Hamasaka *et al*., 2007; McCarthy *et al*., 2011; Tabuchi *et al*., 2018), GABA (Parisky *et al*., 2008; McCarthy *et al*., 2011; Lelito & Shafer, 2012; Gmeiner *et al*., 2013; Liu *et al*., 2014), and acetylcholine (Wegener *et al*., 2004; McCarthy *et al*., 2011; Lelito & Shafer, 2012). In the context of neuropeptidergic regulation, the neuropeptide diuretic hormones 31 (Dh31) and 44 (Dh44) are homologous to the mammalian calcitonin gene-related peptide (CGRP) and corticotropin-releasing factor (CRF), respectively (Lee *et al*., 2023a). Similar to the mammalian CGRP, *Drosophila* Dh31 plays key roles in intestinal peristalsis (Benguettat *et al*., 2018), reproduction (Kurogi *et al*., 2023), memory formation (Lyu *et al*., 2023), sleep regulation (Kunst *et al*., 2014), circadian rhythms (Goda *et al*., 2019), and temperature compensation (Goda *et al*., 2016). The conserved functions of Dh31 across species underscore its importance in the regulation of diverse physiological processes. Similarly, Dh44 and CRF are involved in the regulation of stress responses, but also circadian regulation of sleep. Mammalian CRF neurons in the paraventricular nucleus of the hypothalamus are involved in the regulation of sleep and wakefulness, as they are part of the neural circuits that control wakefulness and are influenced by the circadian pacemaker (Ono *et al*., 2020). In *Drosophila*, Dh44-positive PI neurons have been implicated in the control of rest-activity rhythms (Cavanaugh *et al*., 2014; King *et al*., 2017). Hugin-expressing neurons, which act downstream of clock neurons to regulate rhythmic locomotor activity, are also suggested as a specific neural circuit through which Dh44 influences these behaviors (King *et al*., 2017; Barber *et al*., 2021; Schwarz *et al*., 2021). In addition, like Dh31, Dh44 is involved in several other aspects of physiology, including growth and metabolism (Dus *et al*., 2015). Dh44 is known to activate PI neurons to control the rhythm of activity, and the reduction of sleep can be activated by nutrient starvation (Oh & Suh, 2023). In the context of the involvement of Dh31 and Dh44 in DN1 and PI neurons, the connections between DN1a and Dh44 neurons are identified in the early third-instars stage (Poe *et al*., 2023), and multiple pieces of evidence support that DN1p potentially secretes Dh31 (Kunst *et al*., 2014), which might play an important role in modulating PI neurons. However, the neurophysiological and computational basis of how Dh31 and Dh44 affect PI neuron firing, and the resulting neural coding is not well understood. In the present study, we identify synergistic effects of Dh31 and Dh44 in terms of how they influence membrane potential dynamics of PI neurons. We find that such a synergistic interaction between Dh31 and Dh44 contributes to enhancing the reliability of action potential timing and synaptic potentiation that promotes arousal. Taken together, these results suggest a multiplexed neuropeptide-dependent precision and plasticity of spike timing in circadian output neurons as a neural coding mechanism for the circadian clock that underlies sleep in *Drosophila*.

## 2 MATERIALS AND METHODS

### 2.1 Fly Strains

The *Drosophila* strains utilized in this study were acquired from the Bloomington Drosophila Stock Center (Bloomington, IN, USA), except for *UAS-slob RNAi* obtained from Vienna Drosophila Research Center (VDRC: 100987). To target and visualize the PI neurons, *Dilp2-Gal4* line (a gift from Amita Sehgal) and *UAS-CD4-tdGFP* line (BDSC: 35836) were recombined through standard genetic recombination techniques to enable use for electrophysiological recordings. We established a split-GAL4 driver by recombining *R20A02-AD* (BDSC: 70582) and *R18H11-DBD* (BDSC: 69017), and *UAS-NaChBac* (BDSC: 9466) and *UAS-dTRPA1* (BDSC: 26263) were used to activate Dh31^+^-DN1p cells targeted by *R20A02-AD;R18H11-DBD*. All strains, with the exception *of R20A02-AD;R18H11-DBD*, were outcrossed into the *iso31* (BDSC: 5905) genetic background at least 4 times. The flies were nourished with standard *Drosophila* food comprising molasses, cornmeal, and yeast. Flies were stocked in a 25°C incubator (DR-36VL, Percival Scientific, Perry, IA, United States) following a 12h:12h light:dark cycle and kept at 65% humidity. All experiments involved female flies (5-8 days old) and were conducted at 25 °C, adhering to all pertinent ethical regulations for animal testing and research at Case Western Reserve University.

### 2.2 Cell-Free Protein Synthesis

Dh31 and Dh44 DNA were amplified from their template DNA based on gBlocks Gene Fragments (Integrated DNA Technologies) by using UltraRun LongRange polymerase (206442;QIAGEN). For PCR-based amplification of Dh31, 5’-AAAGCGATCGCATGACAAACCGAT-3’ and 5’-GTTTAAACTTAGACATCGGTCTCGG-3’ were used as primers, and for amplification of Dh44, 5’-AAAGCGATCGCATGATGAAAGCCACA-3’ and 5’-GGGCGGTT TAAACTTAATTAACGTTAT-3’ were used as primers. These corresponded to nucleotides 1 to 351 or 3,781 to 4,132 in the Dh31 sequence and corresponded to nucleotides 1 to 1071 or 5,311 to 6,382 in the Dh44 sequence. Amplification was performed by using the following thermal program: 93°C for 3 min; 35 cycles of 95°C for 30 s, 54°C for 30 s, and 65°C for 10 min; followed by one cycle at 72°C for 10 min. The PCR products were separated by electrophoresis on a 1.5% agarose gel. The DNA was extracted from the agarose gel using the GeneJET Gel Extraction Kit (K0691; Thermo Scientific). The samples were purified to reach sufficient concentrations before proceeding to each step. Purification was performed by ethanol precipitation and Wizard SV Gel and PCR Clean-Up System (A9281; Promega). DNA fragments encoding Dh31 and Dh44 were subcloned into the SgfI and PmeI sites of pF25A ICE T7 Flexi Vector (L1061, Promega) available in the kit of the TnT T7 Insect Cell Extract Protein Expression System (L1101, Promega), according to manufacturer instructions. The identities of the subcloned PCR products were verified by Sanger Sequencing analysis by using primers 5’-CGGATGGCCTTTTTGCG TTTCTA-3’ and 5’-CTTCCTTTCGGGCTTTGT TAG-3’ for Dh31, and 5’-AAAGCGATCGCA TGATGAAAGCCACA-3’ and 5’-GGGCGGT TTAAACT TAATTAACGTTAT-3’ for Dh44. For protein synthesis, the TnT T7 Insect Cell Extract Protein Expression System was used to allow both transcription and translation to occur in a single reaction. TnT T7 ICE Master Mix was mixed with 4µg of plasmid DNA template and brought up to a volume of 50 µl with nuclease-free water. The reactions were incubated at 30°C for 4 hours to allow protein synthesis to occur. Synthesized Dh31 and Dh44 were stored at -80 °C until use. These synthesized Dh31 and Dh44 were used at a final concentration of 10^−6^ M when they were tested alone (i.e., Dh31 or Dh44), and at a final concentration of 5 x 10^−7^ M when they were tested as a cocktail (i.e., Dh31 and Dh44). Selected concentration was based on previous study (Shafer *et al*., 2008). As a control vehicle, TnT T7 ICE Master Mix with Luciferase ICE T7 Control DNA was used.

### 2.3. Membrane-coated glass electrodes

We performed perforated patch-clamp and sharp-electrode electrophysiological recordings from a PI neuron using membrane-coated glass electrodes (Jameson *et al*., 2024). We used a perforated patch clamp in order to acquire action potential firing with current-clamp mode and KCa currents with voltage-clamp mode. On the other hand, we used sharp-electrode electrophysiological recordings to acquire synaptic potentials within axonal regions under tonic hyperpolarization current injection, which is hard to hold with the patch clamp technique. We used membrane-coated glass electrodes to make recording more stable as we found that membrane-coated glass electrodes were helpful for suppressing artifactual signal variability, which is mainly derived from access resistance fluctuations during the recording (Jameson *et al*., 2024). A lipid membrane was created with the use of 7.6 g/L egg yolk lecithin (440154; Sigma-Aldrich) and 2 mM cholesterol (C8667; Sigma-Aldrich) by dissolving with hexane solvent (34859; Sigma-Aldrich) and acetone (270725; Sigma-Aldrich) for one hour at room temperature using an ultrasonication machine, as previously described (Jameson *et al*., 2024). Hexane and acetone were evaporated through pressure injection of inert nitrogen, followed by an additional incubation under the decompression chamber. The hexane/acetone solvent was entirely removed, and the egg yolk lecithin and cholesterol were transferred to a mixture of liquid paraffin/squalene (7/3, v/v), then incubated at 80℃ overnight. The following day, the prepared lipid was promptly utilized for electrode preparation. Patch-electrodes (12 − 20 MΩ) were formed using a Flaming-Brown puller (p-97; Sutter Instrument, CA, USA with thoroughly cleaned borosilicate glass capillaries (OD/ID: 1.2/0.68mm). The electrode tip was refined with a microforge (MF200; World Precision Instruments, FL, USA). Sharp-electrodes (120 − 190 MΩ) were formed using a laser-based micropipette puller (P-2000, Sutter instrument) with thoroughly cleaned quartz glass capillaries (OD/ID: 1.2/0.6mm). These electrodes were coated with the prepared phospholipids using a tip-dip protocol (Jameson *et al*., 2024). Briefly, we initially dipped the electrode tip into an internal electrode solution in a custom-made reservoir, and then the prepared lipid solution was loaded onto the surface of the internal electrode solution. Following the application of the minimum amount (<20 μL) of prepared lipid solution onto the surface of the internal electrode solution, the electrode was vertically lifted using a micromanipulator. The electrode was prepared for electrophysiological recording through standard backfilling of the internal electrode solution by using a microfiller (MF34G MicroFil; World Precision Instruments). The internal pipette solution loaded into the tip contained 102 mM potassium gluconate, 0.085 mM CaCl_2_, 0.94 mM EGTA, 8.5 mM HEPES, 4 mM Mg-ATP, 0.5 mM Na-GTP, 17 mM NaCl, pH7.2 and was utilized for all patch-clamp recording experiments. For sharp electrode intracellular recordings, a 1 M KCl internal pipette solution was loaded into the pipette. Filtering of the internal pipette solutions were performed using a syringe filter with a pore size of 0.02 μm (Anotop 10, Whatman).

### 2.4. Preparation

*In vivo* preparation for electrophysiology of PI neurons was performed as previously described (Tabuchi *et al*., 2018). To anesthetize the flies, they were chilled for 10 min and glued to a 0.025 mm thick metal shim using dental wax or UV light cure adhesives. The cuticle was then peeled off to expose the surface of the brain, and the tethered fly was mounted in a chamber containing *Drosophila* physiological saline (101 mM NaCl, 3 mM KCl, 1 mM CaCl_2_, 4 mM MgCl_2_, 1.25mM NaH_2_PO_4_, 20.7 mM NaHCO_3_, and 5 mM glucose; with osmolarity adjusted to 235-245 mOsm and pH 7.2), which was pre-bubbled with 95% O_2_ and 5% CO_2_. The large trachea and intracranial muscles were removed. The glial sheath surrounding the PI neurons was carefully removed using sharp forceps following the treatment of the enzymatic cocktail, collagenase (0.1 mg/mL), protease XIV (0.2 mg/mL), and dispase (0.3 mg/mL), at 22°C for one minute, which is essentially the same condition when we conduct electrophysiological recordings from DN1 neurons. A small stream of saline was repeatedly pressure-ejected from a large-diameter pipette to clean the surface of the cell body under a dissection microscope. The PI neurons were visualized via tdGFP fluorescence by utilizing the PE4000 CoolLED illumination system (CoolLED Ltd., Andover, UK) on a fixed-stage upright microscope (BX51WI; Olympus, Japan).

### 2.5. Patch-clamp electrophysiology

Perforated patch-clamp was conducted using somatic recording, following procedures described in previous studies (Nguyen *et al*., 2022; Jameson *et al*., 2024). Escin was prepared as a 50 mM stock solution in water and was added fresh to the internal pipette solution, achieving a final concentration of 50 μM. To prevent light-induced changes, the filling syringes were wrapped with aluminum foil due to the light-sensitivity of escin. The escin pipette solutions demonstrated stability for several hours after mixing in the filling syringe, with no observable formation of precipitates. Junction potentials were nullified before the formation of a high-resistance seal formation. Perforated patches were allowed to spontaneously develop over time following the high-resistance seal formation. Once breakthrough was apparent, indicated by the gradual development of a large capacitance transient in the seal test window, monitoring of access resistance was initiated using the membrane test function. From that point onward, access resistance (Ra) was continuously monitored throughout the final stages of the perforation process until it stabilized (Ra stably < 40 MΩ). To isolate KCa currents, we perfused saline containing 10^−7^ M Tetrodotoxin to block voltage-gated sodium currents and 4 × 10^−3^ M 4-Aminopyridine to block fast-inactivated voltage-gated potassium currents (Tabuchi *et al*., 2018). The PI neurons were held at −70 mV and two series of 200-ms voltage pulses were delivered in 10-mV increments between −80 and 60 mV, with a -90mV pre-pulse delivered before each voltage push. The second series was recorded with saline containing 5 × 10^−4^ M CdCl_2_, which abolished voltage-activated Ca^2+^ currents. The subtraction of the current trace in the presence of 5 × 10^−4^ M CdCl_2_ from the current trace without 5 × 10^−4^ M CdCl_2_ was defined as KCa current. Axopatch 1D amplifier (Molecular Devices) was utilized in obtaining electrophysiological signals, which were then sampled with Digidata 1550B (Molecular Devices) under the control of pCLAMP 11 (Molecular Devices). The electrophysiological signals were sampled at 20 kHz and subjected to a low-pass filter at 1 kHz. To characterize Dh31 and Dh44 inducible responses, we perfused apamin (10^−5^ M) and/or cadmium (10^−4^ M) together with Dh31 and Dh44 (Jedema & Grace, 2004).

### 2.6. Sharp electrode intracellular recordings

Intracellular recording was performed by inserting sharp microelectrodes as described in previous studies (Nguyen *et al*., 2022; Jameson *et al*., 2024). We utilized this method in order to acquire stable synaptic potentials under tonic hyperpolarization current injection. The electrode was introduced into the region characterized by dense tdGFP signals within the axonal regions of PIs in *Dilp2-Gal4>UAS-CD4-tdGFP* flies. During the insertion of the electrode, tdGFP signals served as the basis for initial visual inspection, while the depth of insertion was directed by alterations in sound (Model 3300 Audio Monitor, A-M Systems) and the generation of membrane potential. To facilitate this process, “buzz” pulses were added just before the electrode was ready to cross the membrane. The duration of the pulse was determined by a technique called “advance air shooting”. If the microscope revealed that the electrode tip was physically shaking, the duration was considered excessive. We utilized the longest duration in the range where the electrodes remained stationary, typically between 2-5 ms. Commencement of membrane potential recordings occurred once the membrane potential had stabilized, typically requiring several minutes. Recordings were conducted with an Axoclamp 2B with HS-2A x 1 LU headstage (Molecular Devices), and sampled with Digidata 1550B interface, controlled by pCLAMP 11 software on a computer. The signals were sampled at 10 kHz and subjected to a low-pass filter at 1 kHz.

### 2.7. Immunostaining

Brains or thoracic ganglion were fixed in 4% PFA at 4°C overnight. After several washes with phosphate-buffered saline (137 mM NaCl, 2.7 mM KCl, 10 mM Na_2_HPO_4_, 1.7 mM KH_2_PO_4_) + 0.3% Triton X-100 (PBST), samples were incubated with rabbit anti-GFP (Thermo Fisher, 1:200) and mouse anti-BRP (nc82, Developmental Studies Hybridoma Bank, 1:50) at 4°C overnight. After additional PBST washes, samples were incubated with Alexa488 anti-rabbit (Thermo Fisher, 1:1000) for anti-GFP staining and Alexa568 anti-mouse (Thermo Fisher, 1:1000) for anti-BRP stainings overnight at 4°C. After another series of washes in PBST at room temperature over 1 hr, samples were cleared in 70% glycerol in PBS for 5 min at room temperature and then mounted in Vectashield (Vector Labs). Confocal microscope images were taken under 10x or 63x magnification using a Zeiss LSM800.

### 2.8. Sleep behavioral assay

Sleep behavior was measured with the use of consolidated locomotor inactivity, as described previously (Tabuchi *et al*., 2018). Female flies (5-8 days old) were loaded into glass tubes containing 5% sucrose and 2% *Drosophila* agar medium. Fly behavior was monitored using the *Drosophila* activity monitors (DAM, Trikinetics, Waltham, MA) in an incubator at 25°C with a 12 hr:12 hr light:dark (LD) cycle for 2 days to measure sleep. The first day following loading was not included in the analysis. Activity counts were collected in 1 min bins, and sleep was identified as periods of inactivity of at least 5 min. Sleeplab (Joiner *et al*., 2006), a MATLAB-based (MathWorks) software, was used to quantify sleep behavior.

### 2.9. Data analysis and statistics

Statistical analyses were done using Prism software version 10.1.0. (GraphPad), Clampfit version 10.7 (Molecular Devices) or MATLAB 2023b (MathWorks). For comparisons of two groups of normally distributed data, unpaired t-tests were used. Paired t-tests were applied for comparisons of electrophysiological signals with normal distributions from the same neuron, before and after stimulation. For comparisons between multiple groups, either one-way ANOVA with post hoc Tukey test or Kruskal-Wallis test with Dunn’s multiple comparisons test was used for normally distributed and non-normally distributed data, respectively. A significance threshold of p-value < 0.05 denoted statistically significant test results while asterisks indicated varying levels of significance (*p < 0.05, **p < 0.01, ***p < 0.001, and ****p < 0.0001). Error bars represent means ± SEM averaged across experiments.

The reliability of the spike onset was calculated using Cronbach’s alpha, as previously described (Tabuchi *et al*., 2018). Cronbach’s alpha is defined as

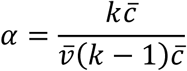

Where *k* represents the number of action potentials, *v̅* represents the average variance of action potential onset rapidness slope, which was defined by dVm/dt measured from spike onset threshold to peak dVm/dt, and *c̅* represents the average inter-dataset covariance of action potential onset rapidness slope.

To quantify spike patterns, the coefficient of variation (CV) and CV2 of interspike intervals (ISI) were used (Holt *et al*., 1996). CV reflects a global measure of irregularity and is defined as

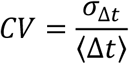

where σis the standard deviation of ISI and μis the average time of ISI.

CV2 indicates a local measure of irregularity defined as the dispersion of the adjacent ISIs. Thus, CV2 is defined as

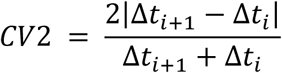

where Δt is an interval time of the *ith* ISI.

We also calculated the local variation (LV) as alternative measures of local irregularity, by computing the dispersion of the two adjacent ISIs. LV is defined as

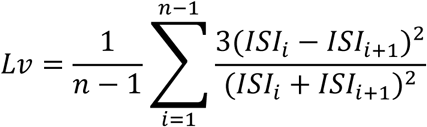

where *ISI_i_* is the *ith* ISI and n is the number of ISIs (Shinomoto *et al*., 2003). While both CV2 and LV compare successive ISIs, they provide different perspectives on spike train variability. CV2 is a measure of overall ISI variability, while LV focuses on the local irregularity of ISI sequences. LV is specifically designed to detect variations in local firing patterns that CV2 may miss, especially in non-Poissonian processes (Shinomoto *et al*., 2009). Although the results for CV2 and LV may appear similar, the inclusion of both metrics is critical for a more complete analysis. CV2 provides insight into general ISI variability, while LV highlights local variations that are important for understanding subtle differences in firing patterns that may not be captured by CV2 alone. Thus, both metrics complement each other to provide a more robust characterization of activity patterns.

To determine the peak firing rate (Hz), the largest 1/ISI value representing the instantaneous frequency for each dataset was used. To quantify the decay time constant (т) for each condition, we used the plot of 1/ISI (Hz) versus time (s) and fit an exponential decay model to the data.

To visualize local PSPs variability, we used a Gaussian Mixture Model (GMM) as described previously (Tabuchi *et al*., 2018). GMM assumes that data points arise from a mixture of Gaussian distributions, each defined by its mean and covariance matrix. Model parameters, including mixture weights and Gaussian component parameters, were estimated using the Expectation-Maximization (EM) algorithm. To preprocess the data, interval timings of PSPs were normalized by the cell-average firing rates and aggregated across cells. Second-order distributions were then constructed by logarithmically binning adjacent pairs of normalized intervals into a 2D histogram. Specifically, we utilized a grid of 22 x 22 bins. For different conditions, distinct logarithmic ranges were employed. We fitted 3 to 5 Gaussian components with full covariance to the joint second-order distributions. For validation purposes, we generated size-matched random samples from the model, producing joint and marginal distributions that closely resembled the training data. To create rate-matched pairs, we first sampled an average PSPs frequency from a beta distribution fitted to the PSPs of PI neurons at ZT18-20. Subsequently, we iteratively constructed novel PSPs trains by selecting the next interval based on the current interval, continuing until the required number of PSPs was reached. Interval selection was performed via rejection sampling of the continuous conditional probability densities of the GMM, following the exclusion of 200 burn-in samples. The resulting logarithmic intervals were exponentiated and normalized to yield PSP times, which were then binned into 10 ms binary signals. To assess the processing performance of the synaptic transmission, we used the area under the curve (AUC) in receiver operating characteristic (ROC) analysis. The AUC in ROC analysis provides a single scalar value that summarizes the performance of a classification model across different thresholds, reflecting the model’s ability to discriminate between different classes, making it a useful metric for assessing processing power in tasks where discrimination is important (Hajian-Tilaki, 2013). A model with an AUC close to 1 indicates strong discriminative power, while an AUC close to 0.5 indicates random guessing (Corbacioglu & Aksel, 2023). Defining performance based on AUC provides a clear and interpretable measure of model effectiveness. We used the 8900 convergence ratios (from δ = 0.01 to δ = 0.9) of presynaptic to postsynaptic transmission as 8900 input arguments to the ROC analysis.

## 3 RESULTS

### 3.1 Dh31/Dh44 induced change in action potential firing

The synapses between DN1 circadian clock neurons and PI neurons play a critical role in translating circadian rhythms into physiological outputs. We used this connection as a model to study multiplexed neuropeptide mechanisms because this synapse is where Dh31 and Dh44 signaling cross. To access this synapse, we used an in vivo preparation in which the postsynaptic PI neurons are genetically labeled (Figure 1a). We focused on ZT18-20 as a specific time window for our electrophysiological recordings because this time window is associated with high-quality sleep during the night, which is supported by a steady state of stable neural dynamics of DN1 circadian clock neurons. PI neurons have less spontaneous firing activity during the night, and ZT18-20 has a particularly low rate of spontaneous firing frequency, which is close to 0 Hz (Tabuchi, 2018). Despite the physical disturbances associated with live dissections of the samples, which is required for our recordings, we find the PI neuron’s clock function is not disrupted. Since both Dh31 and Dh44 are associated with arousal promotion, we hypothesized that bath application of Dh31 and Dh44 would cause an increase in firing activity of PI neurons. We performed cell-free protein synthesis of Dh31 and Dh44 and applied the synthesized Dh31 and Dh44 to PI neurons (Figure 1b). Compared to the control (Figure 1c), spontaneous firing of PI neurons during ZT18-20 increased with the application of Dh31 alone (Figure 1d), Dh44 alone (Figure 1e), and Dh31 and Dh44 together (Figure 1f). We found that a mixture of Dh31 and Dh44 enhanced supralinear firing of PI neurons (Figure 1f). In contrast to the spontaneous firing rate, we did not find significant changes in the resting membrane potential (Figure 1g–j). Next, we quantified intrinsic membrane excitability by evoking activity through current injection (Figure 2a). While the evoked mean firing rate did not increase with either Dh31 or Dh44 alone (Figure 2c-d), the mixture of Dh31 and Dh44 showed a supralinear increase, demonstrating a strong increase in elicited firing frequency (Figure 2e). To further characterize the effects of Dh31 and/or Dh44 applications in PI neuronal intrinsic firing properties, we also analyze peak firing rate and decay time constant. The maximum 1/ISI value, representing instantaneous frequency, was taken as the peak firing rate. The decay time constant (τ) was determined by an exponential decay model fitting to the 1/ISI vs. time plot. Peak instantaneous frequency did not show a significant difference in the control vehicle (Figure 2f) and Dh31 application (Figure 2g), but the Dh44 application showed a significant increase in the peak instantaneous frequency (Figure 2h). Similar to the analysis of spontaneous firing activity (Figure 1) and the mean frequency of the evoked firing (Figure 2e), the mixture of Dh31 and Dh44 showed a more dramatic increase in the peak instantaneous frequency (Figure 2i).

**FIGURE 1.**
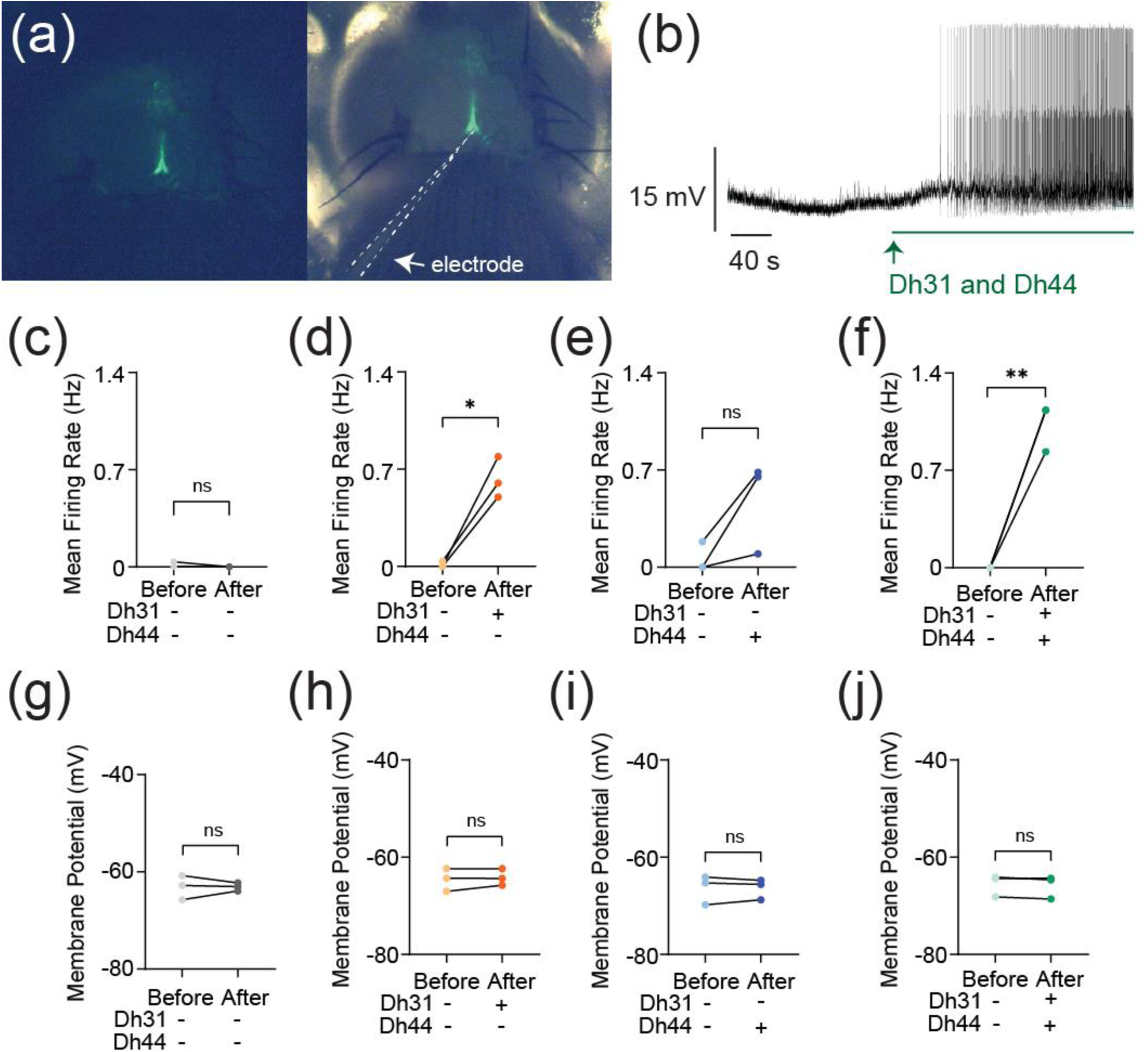
Change in action potential firing induced by Dh31/Dh44. **(a**) Representation of *in vivo* expression of the PI neurons through genetic labeling in the brain of the *Drosophila*. **(b)** Voltage trace during the application of synthesized Dh31 and Dh44 to the PI neurons during patch-clamp recording. **(c-f)** Spontaneous firing of PI neurons during ZT18-20 by using control, with (+) Dh31 (N=3) and without (-) Dh31 (N=3), with (+) Dh44 (N=3) and without (-) Dh44 (N=3) and a combination of Dh31 and Dh44 (N=3). **(g-j)** Resting membrane potential of the PI neurons using control, with (+) Dh31 (N=3) and without (-) Dh31 (N=3), with (+) Dh44 (N=3) and without (-) Dh44 (N=3) and a combination of Dh31 and Dh44. The statistics used were paired t-test with *p < 0.05 and **p < 0.01 and ns indicated non-significant.

**FIGURE 2.**
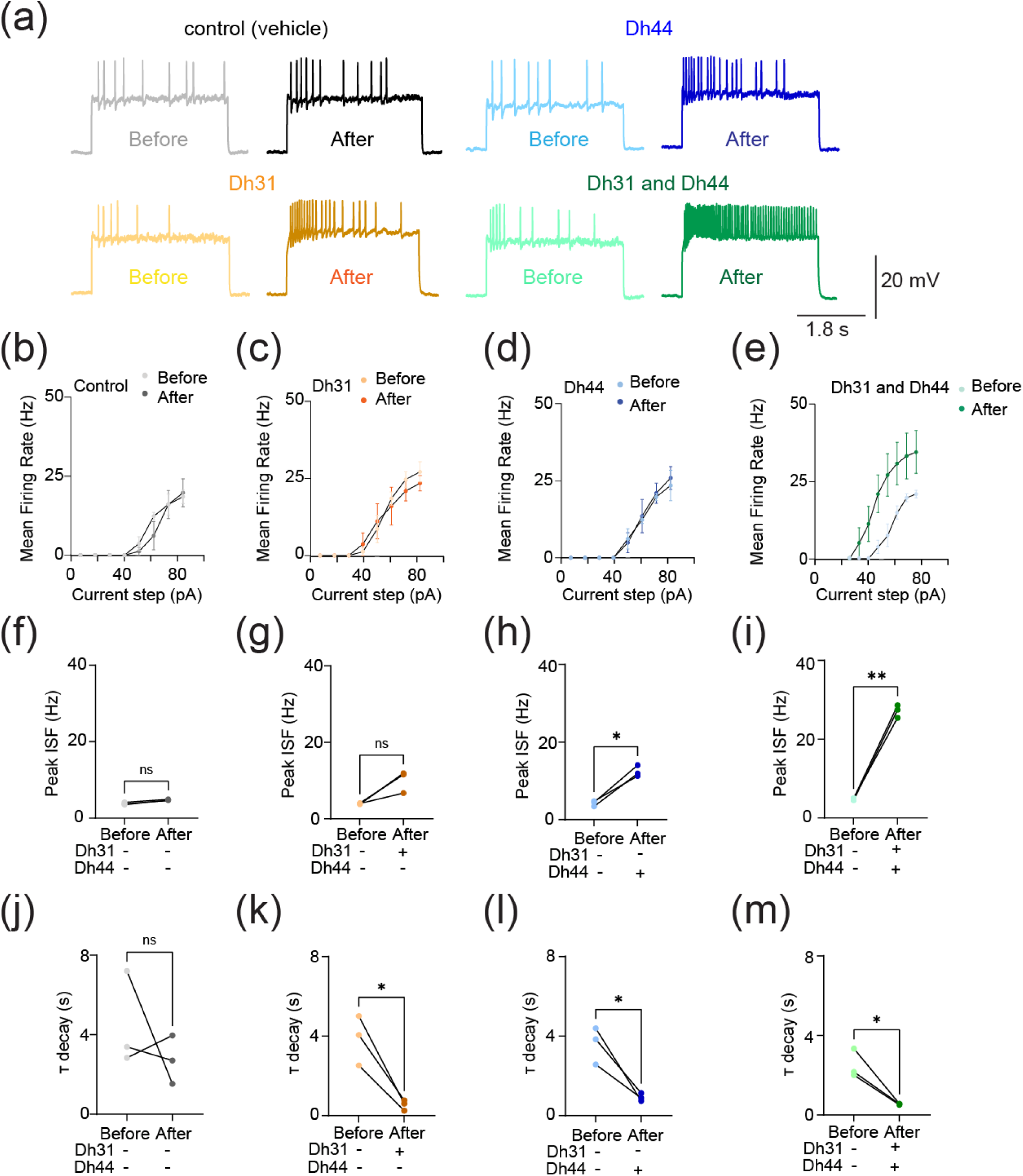
Change in evoked action potential firing induced by Dh31/Dh44. **(a)** Voltage trace before and after the application of synthesized Dh31 and Dh44 to the PI neurons during patch-clamp recording. **(b-e)** Evoked firing frequency by current injection with control, Dh31, Dh44, and a combination of Dh31 and Dh44 (N=3). **(f-i)** Evoked peak firing frequency (Hz) by current injection with control, Dh31, Dh44, and a combination of Dh31 and Dh44 (N=3). **(j-m)** Evoked decay time constant (Τ) for control, Dh31, Dh44, and a combination of Dh31 and Dh44 (N=3). The statistics used were paired t-test with *p < 0.05 and **p < 0.01 and ns indicated non-significant.

The decay time constant, on the other hand, showed a similar level of reduction compared to the control (Figure 2j) in the applications of Dh31 alone (Figure 2k), Dh31 alone (Figure 2l), and the mixture of Dh31 and Dh44 (Figure 2m). Together, these data suggest that Dh31 and Dh44 synergistically interact to enhance the action potential firing of PI neurons during ZT18-20.

### 3.2 Dh31/Dh44 induced action potential waveform change

To investigate the biophysical origins underlying the Dh31/Dh44-induced change in action potential firing, we analyzed the kinetics of the action potential waveform (Figure 3a). We focused on analyzing action potential waveforms from spontaneous firing because the kinetics were artifactually distorted in evoked firing (Figure 2a), which is commonly observed in invertebrate unipolar neurons due to the distance between the current injection site and the action potential generation site (Gouwens & Wilson, 2009). The action potential waveforms exhibit variation induced by interactions with Dh31, Dh44, Dh31/Dh44, Dh31/Dh44 with apamin (10^−5^ M), and Dh31/Dh44 with cadmium (10^−4^ M). Cadmium and apamin were used to selectively inhibit calcium and SK channels, respectively (Jedema & Grace, 2004). The amplitude and threshold of action potentials were not significantly different in the presence of Dh31/Dh44 (Figure 3b, c), but the reduction of amplitude was observed only in the presence of cadmium (Figure 3b), and the increase of threshold was observed only in the presence of cadmium (Figure 3c). A non-significant trend in greater afterhyperpolarization (AHP) amplitude was induced by a mixture of Dh31 and Dh44 (Figure 3d). Interestingly, the change in AHP amplitude was only sensitive to cadmium but not apamin. Based on the potential relationship between action potential waveforms kinetics and firing precision (Axmacher & Miles, 2004; Naundorf *et al*., 2006; Niday & Bean, 2021), we quantified the reliability of action potential timing by using internal consistency of onset rapidness (Tabuchi *et al*., 2018), which we computed using Cronbach’s alpha. We found that the reliability of spike timing was significantly enhanced in the mixture of Dh31 and Dh44, which was reduced by cadmium (Figure 3e). Taken together, these data suggest multifactorial aspects of the biophysical origins underlying the Dh31/Dh44-induced change in action potential firing to make spike timing of PI neurons more reliable in the presence of Dh31 and Dh44 mixture.

**FIGURE 3.**
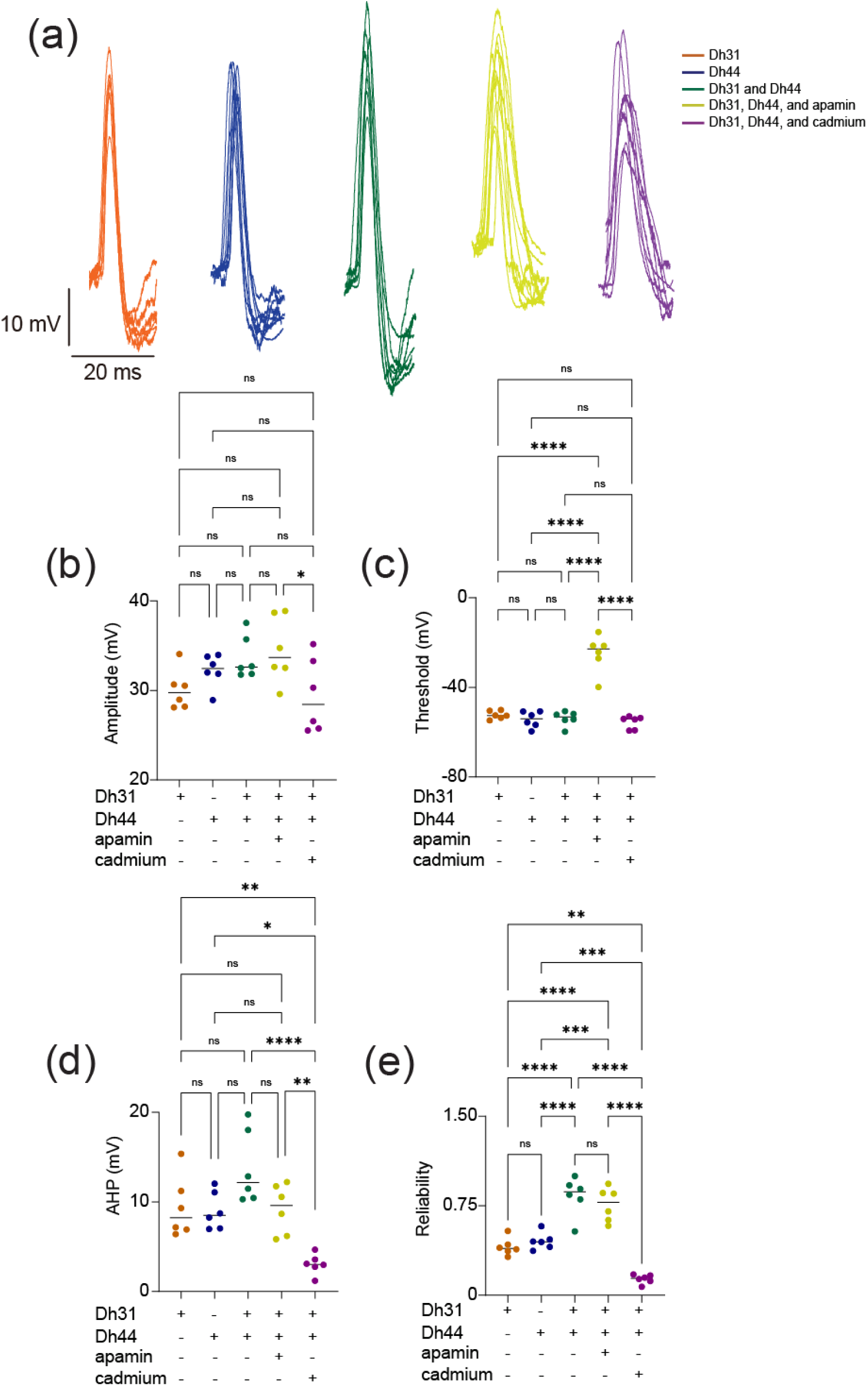
Spontaneous action potential waveform change during the applications of Dh31/Dh44. **(a**) Voltage traces showing action potential waveform changes by Dh31, Dh44, Dh31/Dh44, Dh31/Dh44/apamin, and Dh31/Dh44/cadmium. **(b)** Quantification of action potential amplitude for varying conditions with (+) and without (-) Dh31, Dh44, apamin, and cadmium (N=6 each) **(c)** Quantification of action potential threshold for varying conditions with (+) and without (-) Dh31, Dh44, apamin, and cadmium (N=6 each). **(d)** Afterhyperpolarization (AHP) amplitude for varying conditions with (+) and without (-) Dh31, Dh44, apamin, and cadmium (N=6 each). **(e)** Quantification of reliability for varying conditions with (+) and without (-) Dh31, Dh44, apamin, and cadmium (N=6 each). To compare more than two multiple-group comparisons, one-way ANOVA with multiple comparisons was used. A *p*-value < 0.05 was considered statistically significant, with asterisks indicating *p*-values as follows: \**p* < 0.05, \*\**p* < 0.01, \*\*\**p* < 0.001, and \*\*\*\**p* < 0.0001.

### 3.3 Dh31/Dh44 induced spike pattern change and KCa channel conductance

Spike timing reliability determined by AHP and onset rapidness is closely related to spike patterns in neuronal cells (Scuri *et al*., 2005; Stiefel *et al*., 2013; Dumenieu *et al*., 2015; Trinh *et al*., 2019; Cui *et al*., 2022). To investigate whether Dh31/Dh44 applications also influence spike patterns, we analyzed spike frequency adaptation, which is one of the mechanisms shaping spike patterns. The spike frequency adaptation was quantified from the ratio of the 1st ISI to the nth ISI, and an index value closer to 1 would indicate that no spike adaptation occurred, whereas an index value closer to 0 would indicate significant spike adaptation with prolongation of the ISIs (Figure 4a). The Dh31/Dh44 interact and convey significant spike irregularity compared to Dh31 alone and Dh44 alone (Figure 4b–d). CV (Figure 4b), LV (Figure 4c), and CV2 (Figure 4d) each provide complementary information about the temporal structure of neuronal spike trains: CV is sensitive to changes in the overall firing rate, but LV is a measure of variability in the timing of spikes within local time windows, and CV2 is a measure of spike irregularity that considers pairs of consecutive ISIs. Nevertheless, we found that the application of cadmium and apamin decreased Dh31/Dh44-enhanced spiking irregularity in all metrics (Figure 4b–d). One of the contributing factors in shaping AHP and spike patterns can be KCa channel conductance (Engbers *et al*., 2012; Stiefel *et al*., 2013; Sahu & Turner, 2021), and KCa channels are known to play a critical role in shaping the activity rhythms of PI neurons (Ruiz *et al*., 2021). Therefore, we directly quantified KCa channel conductance in PI neurons by voltage-clamp recordings (Figure 4e). Similar to our previous AHP analysis (Figure 3d), we found that a non-significant trend in greater K_Ca_ conductance amplitude was induced by a mixture of Dh31 and Dh44 (Figure 4 f, g). SLOB (slowpoke-binding protein) is known as a key mediator in modifying K_Ca_ conductance (Shahidullah *et al*., 2009; Jepson *et al*., 2013). Thus, we tested if genetic manipulation of SLOB in PI neurons changes Dh31/Dh44-induced changes in K_Ca_ conductance. Strikingly, we found that Dh31/Dh44-induced changes in K_Ca_ conductance was completely eliminated by introducing SLOB RNAi in PI neurons (Figure 4f, g). These results suggest that Dh31/Dh44-induced changes in K_Ca_ conductance were mediated by SLOB in PI neurons.

**FIGURE 4.**
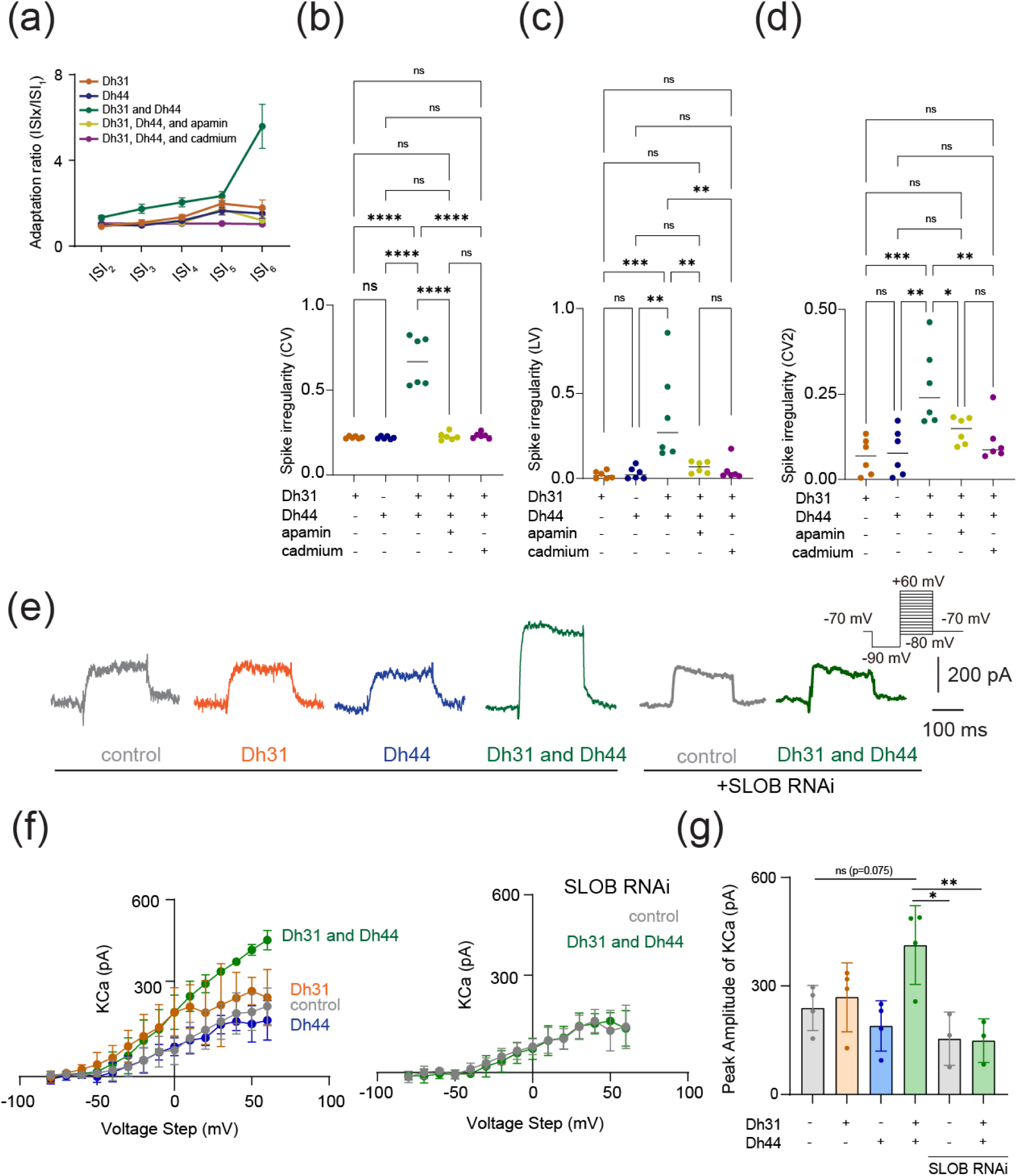
Dh31/Dh44-induced changes in spike pattern and KCa channel conductance. **(a)** Quantification of spike frequency adaptation with ISIx/ISI_1_ ratio for Dh31, Dh44, Dh31 and Dh44, Dh31/Dh44/apamin, and Dh31/Dh44/cadmium (N=5). **(b-d)** Quantification of spike irregularity of CV, LV, and CV2 for varying conditions with (+) and without (-) Dh31, Dh44, apamin, and cadmium (N=6 each). **(e)** Current traces (at +30mV voltage-step) of control, Dh31, Dh44, Dh31/Dh44, SLOB RNAi control, and SLOB RNAi with Dh31/Dh44. **(f)** I-V relationship of KCa conductance and peak amplitude of K_Ca_ for control, Dh31, Dh44, and Dh31/Dh44 (n=4 each), as well as for SLOB RNAi control and SLOB RNAi with Dh31/Dh44 (n=3 each). **(g)** Quantification of Peak Amplitude of KCa for varying conditions with (+) and without (-) Dh31, Dh44 (n=4 each), and SLOB RNAi conditions (n=3 each). Two-way ANOVA with multiple comparisons was used. A *p*-value < 0.05 was considered statistically significant, with asterisks indicating *p*-values as follows: \**p* < 0.05, \*\**p* < 0.01, \*\*\**p* < 0.001, and \*\*\*\**p* < 0.0001.

### 3.4 Dh31/Dh44 induced changes in PSP amplitude

To reveal if Dh31/Dh44 impacted synaptic drive, we examined postsynaptic potential (PSP) parameters such as amplitude, inter-event interval, cumulative probability, and probability density. The voltage traces of PSPs are altered when Dh31 and Dh44 were introduced, with high increased activity when both Dh31/Dh44 were present (Figure 5a). The cumulative probability varied for each condition when recording across PSP amplitudes (Figure 5b). The amplitude of PSP for the condition with Dh31 alone and Dh44 alone was significantly greater compared to the control (Figure 5c). The amplitude of PSP for the condition with both Dh31 and Dh44 was significantly greater than not only the control, but also Dh31 alone and Dh44 alone. (Figure 5c). The addition of apamin in the mixture of Dh31 and Dh44 brought the effects back to the level of Dh31 alone and Dh44 alone, but not to control (Figure 5c).

**FIGURE 5.**
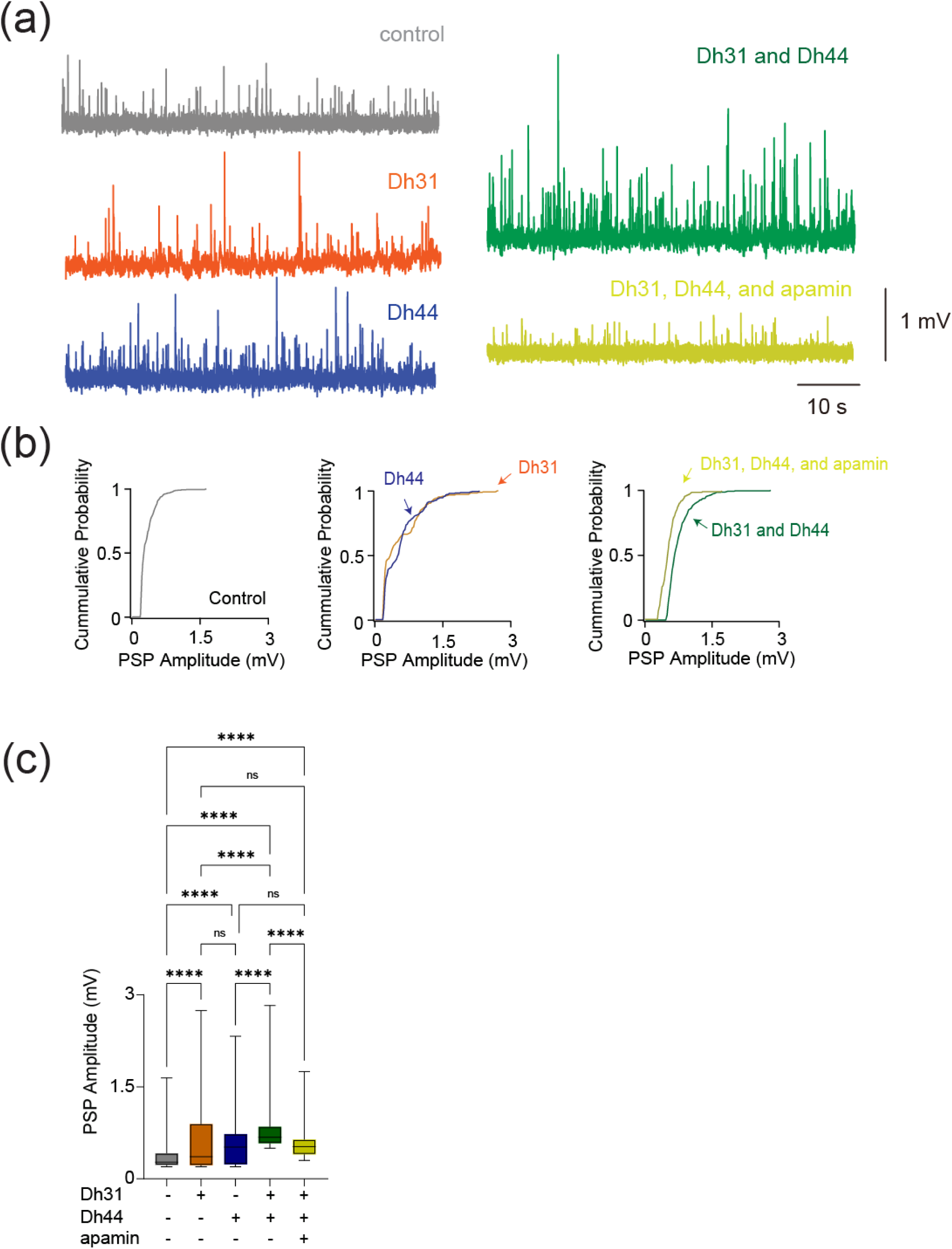
Changes in synaptic potential induced by Dh31/Dh44. **(a**) Voltage traces showing synaptic potential sequence changes for control, Dh31, Dh44, Dh31/Dh44, and Dh31/Dh44/apamin. **(b)** Cumulative probability of PSP for control, Dh31, Dh44, Dh31/Dh44, and Dh31/Dh44/apamin.**(c)** Quantification of postsynaptic potential (PSP) amplitude for varying conditions with (+) and without (-) Dh31, Dh44, and apamin. One-way ANOVA with multiple comparisons was used for the statistics with \*\*\**p* < 0.001, and \*\*\*\**p* < 0.0001.

### 3.5 Dh31/Dh44 induced changes in PSP frequency and temporal patterns of synaptic potentials

Amplitude changes in PSP generally reflect postsynaptic changes in response to neurotransmitters (Turrigiano *et al*., 1998), and we next asked whether the frequency of PSP, which is more related to presynaptic events, was changed. To characterize potential presynaptic changes, the frequency of PSP was analyzed. The probability density during PSP inter-event intervals for the Dh31/Dh44 and Dh31/Dh44/apamin groups was decreased (Figure 6a). By quantifying the PSP inter-event interval for each condition, we found that Dh31/Dh44/apamin was significantly different from all other conditions in that its inter-event interval was longer, indicating a reduction in synaptic activity (Figure 6b). Taken together, these data suggest that Dh31 and Dh44 interact synergistically to enhance the PSPs of PI neurons, which may be mediated by both presynaptic and postsynaptic mechanisms. To identify the computational basis for how Dh31/Dh44 alters the temporal dynamics of synaptic potentials, we implemented two types of computational models. We first used a Gaussian Mixture Model (GMM). Interval timings of two adjacent PSPs were projected into two-dimensional space, and GMM was used to estimate the geometry of their probability distribution (Figure 6c). We found the emergence of an additional cluster in the presence of Dh31 or Dh44 compared to the singular cluster in the control. Furthermore, in the presence of Dh31 and Dh44, we found the emergence of multiple clusters which was partially diminished by the administration of apamin. We next conducted receiver operating characteristic (ROC) to estimate the processing performance of interval timing of postsynaptic activity of PI neurons. The higher the area under the curve (AUC) in the ROC analysis, the higher the discriminative power, while an AUC close to 0.5 indicates a random chance level (Corbacioglu & Aksel, 2023). We found that the interval timing of PSPs in the presence of Dh31 or Dh44 alone showed higher AUC compared with the control, but their AUC was lower than the interval timing of PSPs in the presence of both Dh31 and Dh44 with the effect partially reduced by apamin (Figure 6d).

**FIGURE 6.**
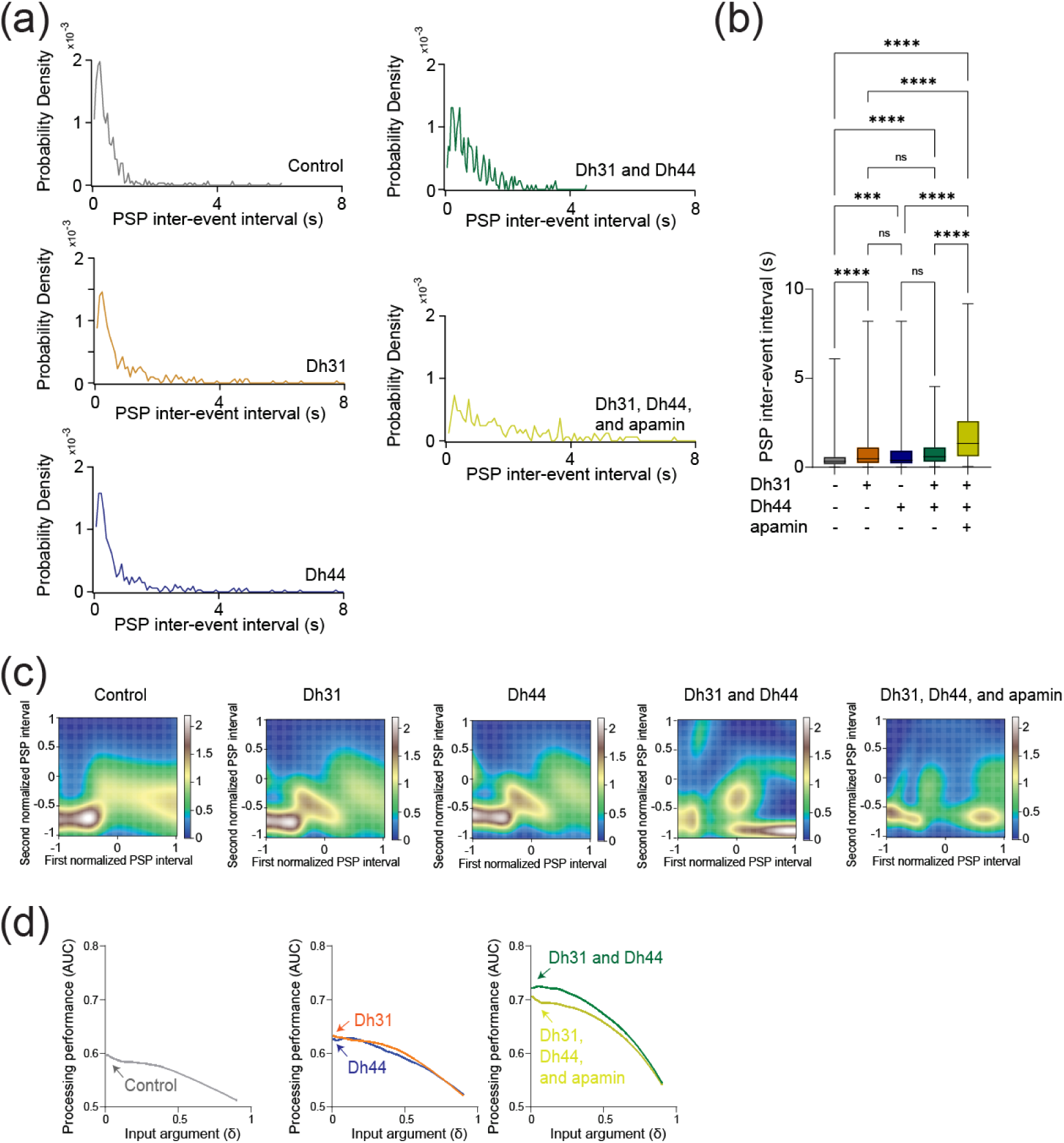
Synaptic potentials show complex patterns under Dh31/Dh44 influence and possible involvement of Dh31-positive DN1 clock neurons. **(a)** Probability density of PSP inter event intervals for control (N=3), Dh31 (N=3), Dh44 (N=30), Dh31/Dh44 (N=3), and Dh31/Dh44/apamin (N=4). **(b)** Time between each event, PSP inter-event interval, was quantified from conditions with (+) and without (-) Dh31, Dh44, and apamin. **(c**) The interval times of two adjacent PSPs were projected into two-dimensional space, and Gaussian Mixture Model (GMM) was used to computationally capture the geometry of their probability distribution. (**d**) The area under the curve (AUC) in receiver operating characteristic (ROC) analysis based on interval timing of PSPs in PI neurons. The statistics used were unpaired t-test with *p < 0.05 and **p < 0.01 and ns indicated non-significant.

### 3.6 Possible involvement of Dh31-positive DN1 clock neurons as demonstrated in sleep alterations

DN1 clock neurons are known to be one of the major presynaptic inputs to PI neurons (Barber *et al*., 2016; Tabuchi *et al*., 2018), and the subset of DN1 clock neurons expressing Dh31 is known to promote wakefulness (Kunst *et al*., 2014). DN1 clock neurons are known to be heterogenous in their role of promoting sleep and/or wakefulness (Guo *et al*., 2018; Lamaze *et al*., 2018), so we decided to establish a split-GAL4 driver to define Dh31-DN1ps neurons. *R20A02-AD* expresses the p65 activation domain associated with Dh31, and *R18H11-DBD* expresses the DNA binding domain of GAL4 associated with PDF receptors (Dionne *et al*., 2018). Thus, we used *R20A02-AD;R18H11-DBD* as an intersectional approach to genetically extract a putative subset of Dh31^+^-DN1p clock neurons. We found that *R20A02-AD;R18H11-DBD>UAS-thGFP* flies label DN1p cells (5.1 cells ± 1.5, N=90 brains) and a few uncharacterized cells in VNC (Figure 7a). To investigate the potential involvement of Dh31-positive DN1 clock neurons, we first assessed whether genetic neural activation in *R20A02-AD;R18H11-DBD* flies would lead to changes in sleep and activity patterns using *UAS-NaChBac*. (Figure 7b). No significant difference in daytime (ZT0-12) sleep was found, but a significant decrease in nighttime (ZT12-24) sleep was found in flies expressing *UAS-NaChBac* in putative Dh31-DN1ps (Figure 7c). We also found an increase in sleep bout number during nighttime in flies expressing *UAS-NaChBac* in putative Dh31-DN1ps (Figure 7d). We also analyzed locomotor profiles (Figure 7e) and found increases in both nighttime activity time (Figure 7f) and activity level (Figure 7g) were observed in flies expressing *UAS-NaChBac* in putative Dh31-DN1ps; it is possible that the expression of NaChBac drove hyperactivity. Constitutive expression of *UAS-NaChBac* can cause artifactual effects not only during experiments but also during development. To exclude this possibility, we also performed acute activation of Dh31-DN1ps based on Thermogenetics. As a thermogenic tool, we used dTRPA1, a temperature-sensitive ion channel, which can be activated by raising the temperature (Hamada *et al*., 2008). We found that activation of putative Dh31-DN1ps neurons by temperature elevation significantly reduced sleep of *R20A02-AD;R18H11-DBD>UAS-dTRPA1* flies (Figure 7h), both during the day and at night (Figure 7i), and facilitated sleep fragmentation as evidenced by an increase in the number of sleep bouts (Figure 7j). We also analyzed activity profiles (Figure 7k) and found an increase in active time during nighttime (Figure 7l), but we found daily activity level during wakefulness was reduced, which was opposed to chronic activation of putative Dh31-DN1ps by expressing NaChBac (Figure 7m). We also quantified activity and sleep profiles of flies having NaChBac alone, without *R20A02-AD;R18H11-DBD*, but we did not find any significant difference compared to control (Figure S1). We also performed another thermogenetics experiments to compare *R20A02-AD;R18H11-DBD>UAS-dTRPA1* flies with *UAS-dTRPA1* alone control flies, and found *R20A02-AD;R18H11-DBD>UAS-dTRPA1* flies showed reduction of sleep and increase of activity under temperature elevation (Figure S2), which is similar to results based on the comparison between *R20A02-AD;R18H11-DBD>UAS-dTRPA1* flies and *R20A02-AD;R18H11-DBD>iso 31* control flies (Figure 7 h-m).

**FIGURE 7.**
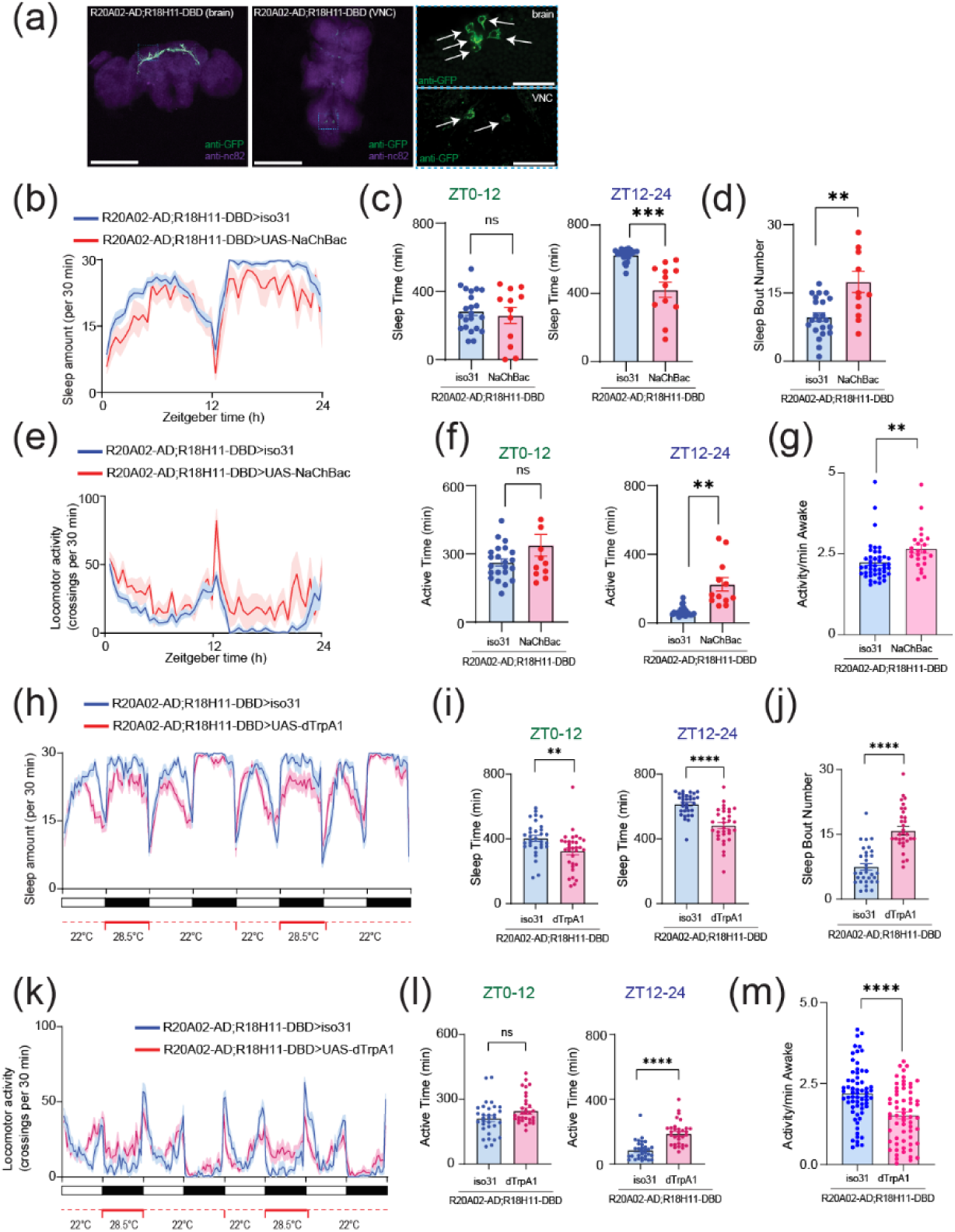
**(a)** Whole-mount brain (left) and thoracic ganglion (right) immunostaining of a *R20A02-AD;R18H11-DBD>UAS-thGFP* fly with anti-GFP (green) and anti-nc82 (magenta). Scale bar indicates 300 μm and 30 μm in inset. **(b)** Sleep profiles **(c)** Sleep time at ZT0-12 and ZT12-24 **(d)** Sleep bout number of *R20A02-AD;R18H11-DBD>iso31* (blue, N=23) and *R20A02-AD;R18H11-DBD>UAS-NaChBac* (red, N=12) flies. Sleep time plotted in 30 min bins. (**e**) Activity profiles (**f**) Active time at ZT0-12 and ZT12-24 (**g**) Daily waking activity of *R20A02-AD;R18H11-DBD>iso31* (blue) and *R20A02-AD;R18H11-DBD>UAS-NaChBac* (red) flies. (**h**) Sleep profiles (**i**) Sleep time at ZT0-12 and ZT12-24 (**j**) Sleep bout number of *R20A02-AD;R18H11-DBD>iso31* (blue) and *R20A02-AD;R18H11-DBD>UAS-dTRPA1* (red) flies. (**k**) Activity profiles (**l**) Active time at ZT0-12 and ZT12-24 (**m**) Daily waking activity of *R20A02-AD;R18H11-DBD>iso31* (blue) and *R20A02-AD;R18H11-DBD>UAS-dTRPA1* (red) flies. The statistics used were unpaired t-test or welch’s t-test with *p < 0.05, **p < 0.01, ***p < 0.001, ****p < 0.0001 and ns indicated non-significant.

## 4 DISCUSSION

The synapses between circadian clock neurons and circadian output cells play a crucial function in translating circadian rhythms into physiological outcomes, regulating internal drives like sleep and hunger. While many molecules that act as output pathways of the circadian clock have been identified, the mechanistic process of how these molecules alter the actual physiology of the network that generates rhythmic internal drives has remained elusive. We show that two types of neuropeptides synergistically interact to modulate circadian clock neural coding. The effects of Dh31 and Dh44 were characterized in experimentation involving both spontaneous (Figure 1) and evoked firing (Figure 2). The induced change in action potential firing in PI neurons from ZT18-20 should indicate an intrinsic membrane excitation change caused by Dh31 and Dh44. We found effects only on firing rate (Figure 1c-f), but not on resting membrane potential (Figure 1g-j). This may indicate that their synergistic mechanisms do not directly come from voltage-dependent conductance, which made us infer that implementing a supplementary pharmacological approach would prove advantageous. Thus, we show that biophysical parameters of the action potential waveform estimate which channels are involved by using apamin and cadmium. The reduction in action potential amplitude was observed only in cadmium treatment, and the increase in threshold were observed specifically in the presence of apamin, suggesting that these effects may involve multifactorial biophysics possibly based on cadmium-sensitive high-threshold calcium channels, and apamin-sensitive channels, including SK channels. Additionally, we observed a non-significant trend toward greater afterhyperpolarization (AHP) amplitude induced by Dh31/Dh44, which was sensitive to cadmium but not to apamin (Figure 3d). This points to a potential role for non-SK potassium (KCa) channels in mediating these effects. In contrast, in Figure 4, both cadmium and apamin blocked Dh31/Dh44 effects on spike adaptation and irregularity. This suggests that while some effects of Dh31/Dh44 signaling involve SK channels (sensitive to apamin), others depend on cadmium-sensitive but apamin-insensitive pathways. Taken together, these findings suggest a complex interplay of ion channels modulated by Dh31/Dh44, potentially including SK channels and other voltage-gated channels. Previous research has shown that apamin specifically blocks SK channels in both *Drosophila* (Abou Tayoun *et al*., 2011) and mammals (Hallworth *et al*., 2003), while cadmium has a broader blocking effect of KCa channels and voltage-gated calcium channels (Jedema & Grace, 2004; Tabuchi *et al*., 2018). From our data (Figure 3), we find that only cadmium has a significant inhibitory effect on neuropeptides, indicating that KCa channels other than SK channels are likely to be the primary effectors. This change suggests that calcium-activated signaling is effective in accelerating repolarization, making the group of neurons susceptible to repetitive firing. We propose that a greater AHP amplitude more effectively resets the membrane potential, leading to quicker recovery and potentially enabling a higher frequency of action potentials. In fact, similar findings have been reported in the mammalian SCN (Cloues & Sather, 2003). We found that greater AHP amplitude induced a change in the availability of voltage-gated conductance for action potential generation, which may be related to a possible mechanism that explains the observed change in amplitude and threshold, suggesting that all the observed changes in action potential waveform kinetics may be related. Because SLOB is known to modulate KCa in both DN1 (Tabuchi *et al*., 2018) and PI neurons (Shahidullah *et al*., 2009; Jepson *et al*., 2013), it is possible that SLOB might be associated with Dh31/Dh44 signaling, and we assess this possibility by directly measuring KCa currents, which have functions that are related to circadian outputs (Ruiz *et al*., 2021) and critical period (Lowe *et al*., 2024).

We next demonstrate that the reliability of the action potential of PI neurons is enhanced by the presence of Dh31 and Dh44 (Figure 3e), and spike irregularity is also enhanced by the presence of Dh31 and Dh44 (Figure 4a-d). The simultaneous induction of increased “reliability” and “irregularity” may appear contradictory. Although neuronal firing is often characterized by irregularity, this doesn’t necessarily mean that it is random (Kostal & Lansky, 2007; Kostal *et al*., 2007; Brette, 2015). Instead, this variability in firing rates can encode important information and reflect the complex dynamics of neural circuits (Kostal & Marsalek, 2010; Waschke *et al*., 2017; Tomar & Kostal, 2021). Our other observation of spike frequency adaptation of PI spiking induced by Dh31 and Dh44 (Figure 4a) further supports this assertion, as spike frequency adaptation is known to enhance the processing performance of information (Benda *et al*., 2005; Salaj *et al*., 2021; Lee *et al*., 2023b) by making the brain not only processes information effectively, but in a way that maximizes the impact of its output on perception and behavior. If irregularity does not necessarily imply randomness, but rather “structured irregularity”, then it is possible that increased reliability is essential for the formation of such structured irregularity (Shinomoto *et al*., 2009; Yang *et al*., 2017; Tabuchi *et al*., 2021; Tabuchi, 2024), which is the idea that the irregularity of neural activity patterns is precisely structured rather than being based on randomness (Chini *et al*., 2022). Interestingly, while spontaneous firing showed an increased decay (Figure 4a), reflected in enhanced spike adaptation, evoked firing exhibited a reduction in decay (Figure 2j-m). The probable explanation for this discrepancy is likely attributable to different mechanisms underlying excitation and firing in these two methods of recording. Invertebrate unipolar neurons, with distinct morphology in comparison to mammalian neurons, initiate action potentials at a greater distance from the soma, such that somatic electrodes cannot accurately control the voltage throughout the cell (Gouwens & Wilson, 2009). Because spontaneous firing utilized the typical dendritic channels to propagate signals and demonstrated a decrease in decay, it is likely a more accurate predictor of the effects of Dh31/Dh44 *in vivo*.

In addition to altering action potentials, Dh31 and Dh44 also alter postsynaptic potentials (PSPs) in the PI neurons. We find that both the amplitude (Figure 5) and frequency (Figure 6) of the PSP epoch were altered by Dh31 and Dh44, suggesting their impacts on both presynaptic and postsynaptic effects. Although retrograde synaptic signaling in the *Drosophila* central synapse has not been thoroughly investigated, it is possible that there is retrograde synaptic signaling related to Dh31 and Dh44 in DN1-PI synapses, as in mammalian synapses (Maejima *et al*., 2001; Wilson & Nicoll, 2001). We found that the changes in PSPs induced by Dh31/Dh44 in PI neurons were sensitive to apamin. Specifically, the Dh31/Dh44-induced increase in PSP amplitude was attenuated in the presence of apamin. Furthermore, apamin increased the PSP inter-event interval, indicating a reduction in neuronal activity. These findings demonstrate that apamin, an SK channel blocker, diminishes synaptic activity properties, potentially through a transsynaptic mechanism mediated by SK channel-dependent signaling. We also find that PSPs are not only significantly enhanced in both amplitude and frequency, but also show more complex temporal structures (Figure 6c). Through the GMM-based visualization of these temporal neural activity patterns (Figure 6c), we can highlight the multidimensional nature in the geometry of neural computation. Theoretical predictions support that such an enhancement is computationally powerful for the precise encoding of information (Levakova *et al*., 2016), and our ROC computational models support this as well (Figure 6d). Even if such theoretical predictions were true, would it be worthwhile to improve the accuracy of neural operations in such a way on millisecond time scales, especially in systems that do not perform fast information processing (such as sensory input), such as circadian clock neurons? Circadian clock neurons must accurately track and represent time at various scales, ranging from milliseconds to hours. Precise encoding across shorter time intervals allows the neurons to cumulatively aggregate these intervals into longer periods, including hours. This accumulation of precise coding spanning several time scales ultimately may contribute to the regulation of sleep/wake cycles by fractal patterns that persist across these hierarchical scales (Hausdorff & Peng, 1996). This hierarchical organization in time perception and regulation may also extend to the internal computations performed by the brain, wherein neuropeptides play a pivotal role in shaping these dynamic processes (Wu *et al*.; Mountoufaris *et al*., 2024).Our results in the present study also support this idea and explain the computational basis of internal state representations based on neural dynamics shaped by multiple neuropeptides. These features of circadian neural coding induced by Dh31 and Dh44 may be behaviorally advantageous for triggering rapid sleep-wake state transitions, particularly when switching from a stable sleep state during the night to a wake state by promoting arousal.

There are several potential limitations to this study. First, we still do not know if Dh31-DN1ps play an important role in Dh31/Dh44 clock neural coding. It is possible that Dh31, deriving from other neurons, plays dominant functions (Goda *et al*., 2019). The same applies to Dh44 in this context. Because PI neurons used in this study are based on *Dilp2-Gal4* line, other types of PI neurons might be responsible for releasing Dh44 (Ruiz *et al*., 2021). If this is the case, Dh44 mechanisms might be based on paracrine signaling rather than autoregulation. Second, it is possible that Dh31/Dh44 signaling is not directly related to the synapse between DN1 and PI neurons, but rather indirectly affected via networks such as Hugin^+^ neurons and/or LNd cells (King *et al*., 2017; Barber *et al*., 2021; Schwarz *et al*., 2021). Addressing these issues may provide more valuable insight into mechanistic frameworks on how Dh31/Dh44 signaling is influencing circadian clock neural coding mechanisms. Taken together, our results demonstrate multiplexed neuropeptide-dependent spike time precision and plasticity as a potential neural coding mechanism of the circadian clock underlying sleep in *Drosophila*.

## DATA AVAILABILITY STATEMENT

The data that support the findings of this study are available from the corresponding author upon reasonable request.

## AUTHOR CONTRIBUTIONS

MT designed the study. BC, VK, DLN, LKS, EMP, ANH, and MT performed the experiments. BC, VK, DLN, MAH, FSF, LKS, EMP, ANH, and MT conducted data analysis. BC and MT wrote the manuscript with input from VK, DLN, MAH, FSF, LKS, EMP, and ANH.

## ACKNOWLEDGEMENTS

We thank Keisuke Sakurai for the loan of electrophysiological equipment, and Joseph Monaco for technical advice on computational modeling. We also thank Ben Strowbridge, Heather Broihier, and Dominique Durand, along with members of the Tabuchi lab for discussion, and the Light Microscopy Imaging Core at Case Western Reserve University for help with confocal microscopy.

## Abbreviations

AHP: afterhyperpolarization
AUC: Area under the curve
CGRP: Calcitonin gene-related peptide
CRF: Corticotropin-releasing factor
CV: coefficient of variation
Dh31: Diuretic hormone 31
Dh44: Diuretic hormone 44
Dilp2: Drosophila insulin-like peptide 2
DN1: Dorsal neuron 1
GMM: Gaussian mixture model
LV: local variation
PCR: Polymerase chain reaction
PDF: pigment-dispersing factor
PFA: paraformaldehyde
PI: Pars Intercerebralis
ROC: receiver operating characteristic
SLOB: slowpoke-binding protein
tdGFP: tandem dimer green fluorescent protein
ZT: zeitgeber time

---

**Figure S1.**
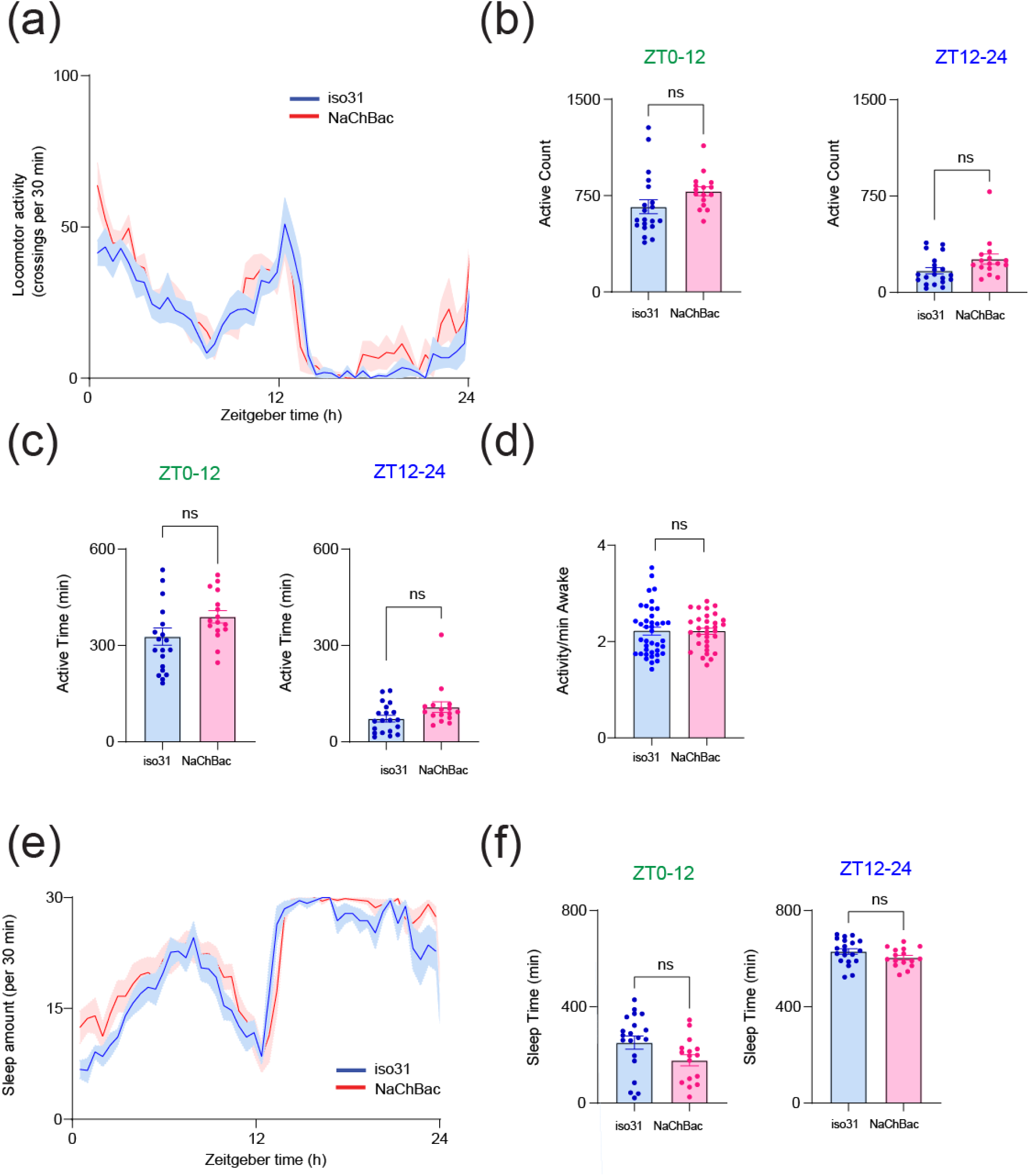
(**a**) Activity profiles (**b**) Activity count at ZT0-12 and ZT12-24 **(c)** Active time at ZT0-12 and ZT12-24 (**d**) Daily waking activity of *iso31* (blue) and *UAS-NaChBac* alone control (red) flies. **(e)** Sleep profiles **(f)** Sleep time at ZT0-12 and ZT12-24 of *iso31* (blue) and *UAS-NaChBac* alone control (red) flies. Sleep time plotted in 30 min bins. The statistics used were unpaired t-tests, and ns indicated non-significant.

**Figure S2.**
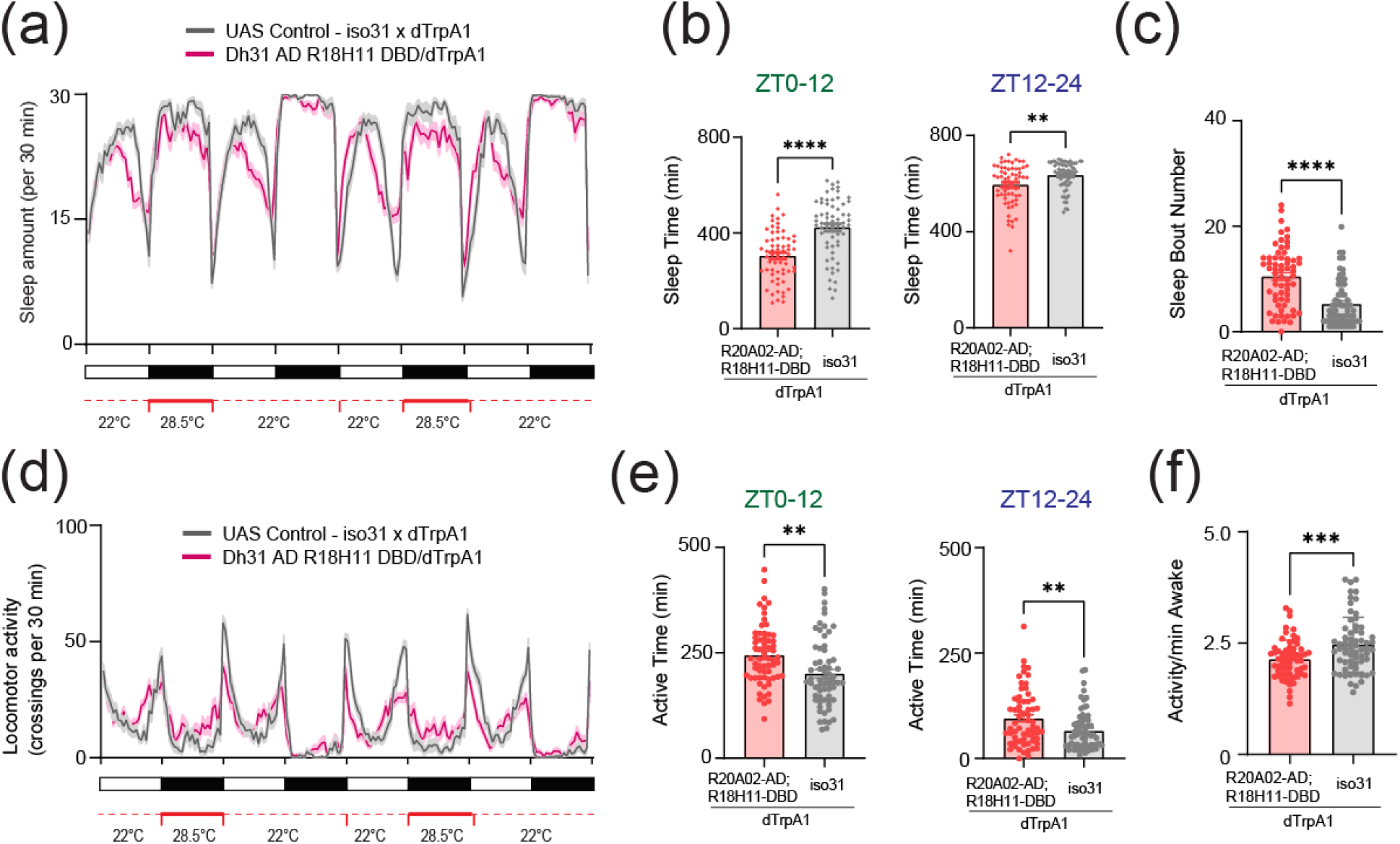
(**a**) Sleep profiles (**b**) Sleep time at ZT0-12 and ZT12-24 (**c**) Sleep bout number of *UAS-dTRPA1>iso31* control (gray) and *R20A02-AD;R18H11-DBD>UAS-dTRPA1* (red) flies. (**d**) Activity profiles (**e**) Active time at ZT0-12 and ZT12-24 (**f**) Daily waking activity of *UAS-dTRPA1>iso31* control (gray) and *R20A02-AD;R18H11-DBD>UAS-dTRPA1* (red) flies. The statistics used were unpaired t-test with **p < 0.01, ***p < 0.001 and ****p < 0.0001.

